# Boolean Logic Coupled with Overrepresentation Analysis Reveals Activation of the Platelet-derived Growth Factor Receptor Beta Pathway in a Model of Oliguric Acute Kidney Injury; Implications for Transition to Chronic Kidney Disease

**DOI:** 10.1101/2025.10.06.680804

**Authors:** Hiral M. Chavre, Terrence Bissoondial, Mahesh Narayan, Prakash Narayan

**Author notes:** Correspondence – Prakash Narayan, PhD, Nodes and Edges LLC, 4030 Wake Forest Rd, Ste 349, Raleigh, NC 27609, USA.

## Abstract

Acute Kidney Injury (AKI) can occur secondary to insults including sepsis, ischemia and contrast dye administration. A time-sensitive increase in serum creatinine (SCr) or reduction in urine output (UO) has been used to define AKI and stage its severity. Oliguria or significantly reduced UO in AKI or oliguric AKI can have a major impact on outcomes including a transition to chronic kidney disease (CKD). We used Boolean logic coupled with overrepresentation analysis to identify the pathway activation signature associated with oliguric AKI in a published study of rat kidney-ischemia reperfusion. In the reperfused kidney, bulk transcriptomic analysis revealed 1068 differentially expressed genes (DEGs). Those DEGs that correlated with UO *and* SCr were submitted to gene ontology biological process overexpression analysis. The pathway activation signature associated with oliguric AKI included positive regulation of profibrotic platelet-derived growth factor receptor beta signaling (fold-enrichment >44) driven by *src, hip1 and hip1r*. Together these findings not only suggest that oliguric AKI may be associated with activation of a pathway leading to fibrosis and CKD but also informs an array of targets to potentially mitigate transition to CKD.

**Highlights:** Mechanistic insights should illuminate therapies. A model of rat kidney-ischemia reperfusion injury was queried by correlating kidney transcriptomics with kidney function to identify the pathway activation signature of oliguric AKI. The most striking feature associated with oliguric AKI was positive regulation of platelet-derived growth factor receptor ß driven by *src, hip1*, and *hip1r*. The pathway activation signature in oliguric AKI informs not only the sequel to injury but also an array of targets to mitigate the potential transition to kidney fibrosis and CKD.

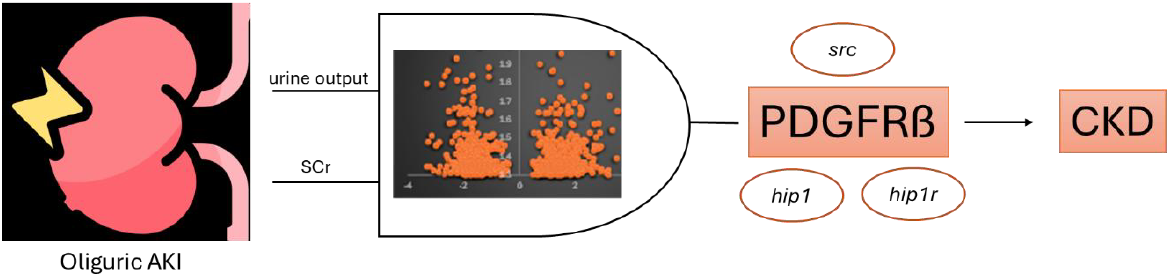

## Introduction

Acute kidney injury (AKI) is a rapid decrease in kidney function evidenced by a time-sensitive increase in serum creatinine (SCr) and/or a reduction in urine output (UO)^1^. Global annual incidence of AKI ∼13.3 million^2^ with several cases occurring in an in-hospital setting. Indeed, AKI can occur secondary to hospital-acquired infection, sepsis, kidney ischemia-reperfusion accompanying complex cardiac surgery and kidney transplant, and following administration of contrast-enhancing dyes^3^. Outcomes in AKI can vary from spontaneous resolution with hydration, to transient requirement for renal replacement therapy (RRT) to death^4-7^. Studies have also reported that survivors of AKI can experience chronic kidney disease (CKD)^8,9^ or an AKI to CKD transition. A severe reduction in UO accompanying AKI, termed oliguric AKI,^10^ can have independent and distinct effects on outcomes vs. non-oliguric AKI. Results from a large study^11^ in an intensive care unit cohort demonstrated that oliguric AKI was associated with 90-day mortality irrespective of concomitant changes in SCr relative to baseline. Intraoperative oliguria was associated with increased in-hospital mortality and need for RRT^12^. In patients with COVID-19-related AKI, oliguria was a predictor of 28-day mortality^13^.

In the present study we analyzed UO and kidney bulk transcriptomic data from a published^14,15^ model of kidney ischemia-reperfusion injury. We used a Boolean logic coupled with overrepresentation analysis to delineate the pathway activation signature accompanying oliguric AKI.

## Methods

Data were analyzed from a previously published study^14,15^ of 45 min unilateral rat kidney ischemia and reperfusion with removal of the contralateral kidney. Kidney bulk mRNA, SCr and UO data were from the control/sham (n=3) and vehicle-treated kidney ischemia-reperfusion (oliguric AKI, n=4) cohorts. Data for UO pertained to 24-hour urine collection levels starting from the onset of reperfusion and data for SCr pertained to measurements made at 24 hours of kidney reperfusion. Bulk mRNA seq data were obtained from control/sham and 24-hour reperfused kidney digests.

All analysis was conducted using MS-Excel. Normalized counts were analyzed following elimination of duplicates and lncRNA (<1% of all counts, data not shown). Differentially expressed genes (DEGs) were identified made using a two-tailed T test followed by the Benjamini–Hochberg test (p_adj_) for multiple comparisons. Determination of the log_2_(fold-change) (FC) was calculated using the formula = log2(average oliguric AKI level/average control). All correlations plots, including the volcano plot, a correlation between log_2_FC and -log10(p_adj_ for p<0.05), were visualized using MS-excel. Correlations between UO and DEGs, were fitted using the equations

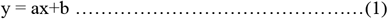

or

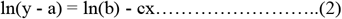

The correlation was considered significant for p<0.01.

The heat map and the t-distributed stochastic neighbor embedding (t-SNE) were visualized using iDEP^16^. Gene Ontology (GO) biological process overrepresentation analysis^17^ was used to determine the pathway activation signature in oliguric AKI. HumanBase^18^ was used to visualize interaction networks between genes in a pathway, and expression of these genes in the kidney.

## Results

Forty-five minutes of kidney ischemia-24 hours reperfusion was associated with a multi-fold reduction in 24-hr UO (Figure 1B) accompanied by a multifold-increase in SCr (Figure 1A) indicative of oliguric AKI.

**Figure 1.**
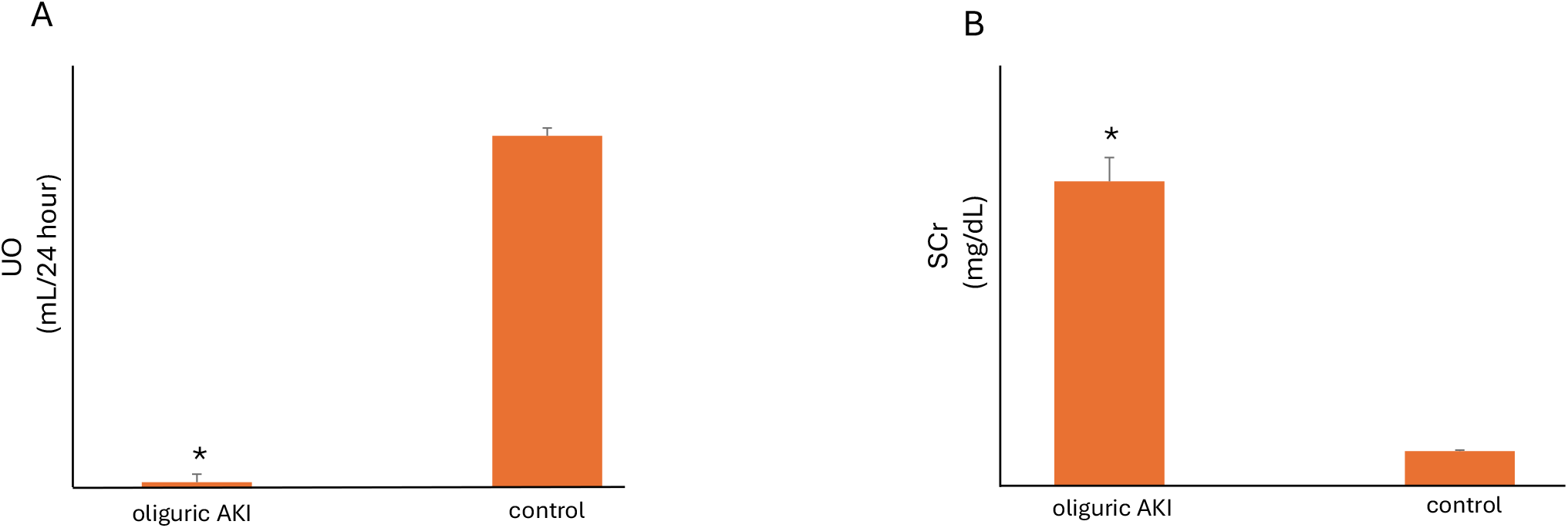
Model of Oliguric AKI. Unilateral rat kidney 45 min ischemia followed by 24 hour reperfusion with removal of the contralateral kidney at the onset of reperfusion is associated with severely reduced UO (A) and an increase in SCr (B) vs. the sham/control cohort. ^*^, p<0.01 vs. control.

This model was associated with 1068 DEGs distributed between 576 upregulated genes and 492 downregulated genes (Figure 2A). Upregulated genes included *havcr1* (168-fold increase), which codes for kidney injury molecule-1 (KIM-1), and *lcn2 (*38-fold increase), which codes for neutrophil gelatinase-associated lipocalin (NGAL)^19^. Heatmap (Figure 2B) and t-SNE (Figure 2C) demonstrated the transcriptomic demarcation between the oliguric AKI and control cohorts.

**Figure 2.**
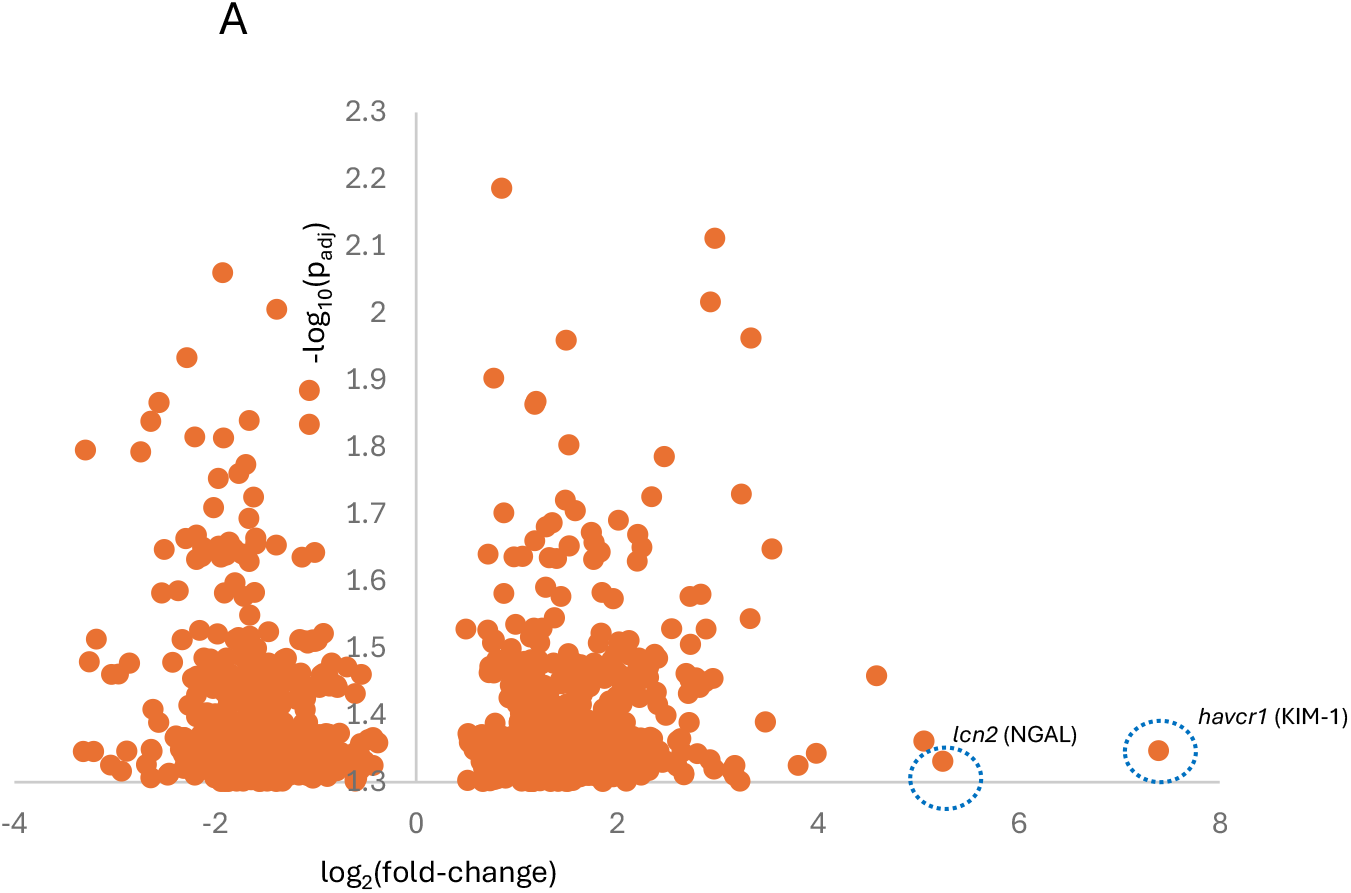
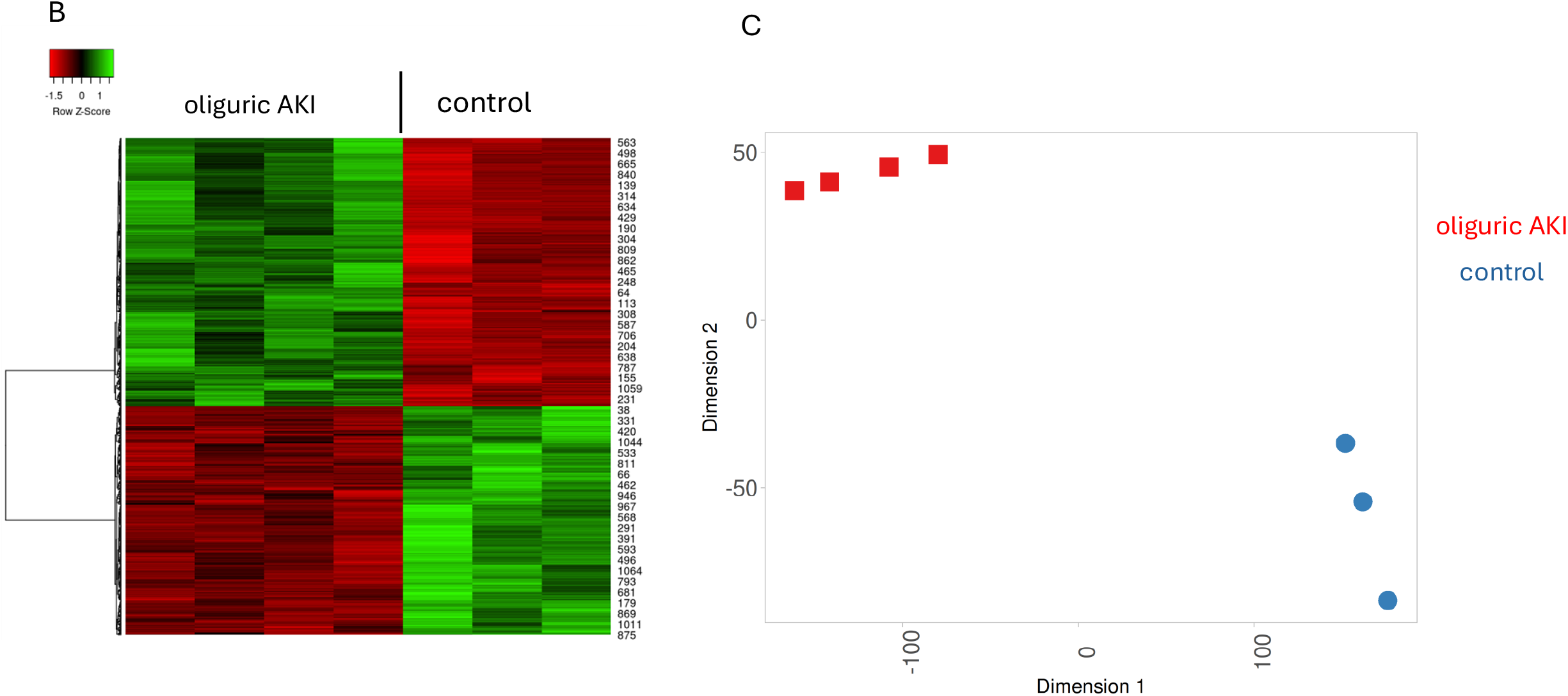
The Oliguric AKI Transcriptome. (A) DEGs (kidney mRNA) with upregulation of *hvacr1* and *lcn2* which code for KIM-1 and NGAL, respectively. Heat map (B) and t-SNE (C) plot showing separation between the oliguric AKI and sham cohorts.

To determine the pathway activation signature associated with oliguric AKI we used a Boolean-gated approach coupled with overrepresentation analysis to identify DEGs that correlated with both UO (Figure 3A) *and* SCr (Figure 3B). Jaccard-Tanimoto similarity analysis^20^ revealed that there is 95.1% similarity between downregulated genes correlating with both UO and SCr, and 95.5% similarity between upregulated genes (Figure 3C) correlating with both UO and SCr (Figure 3D). Nevertheless, the strength of the association (r) between DEGs correlating with UO (Figure 3E) was dissimilar to the strength of the association for DEGs correlating with SCr (Figure 3F).

**Figure 3.**
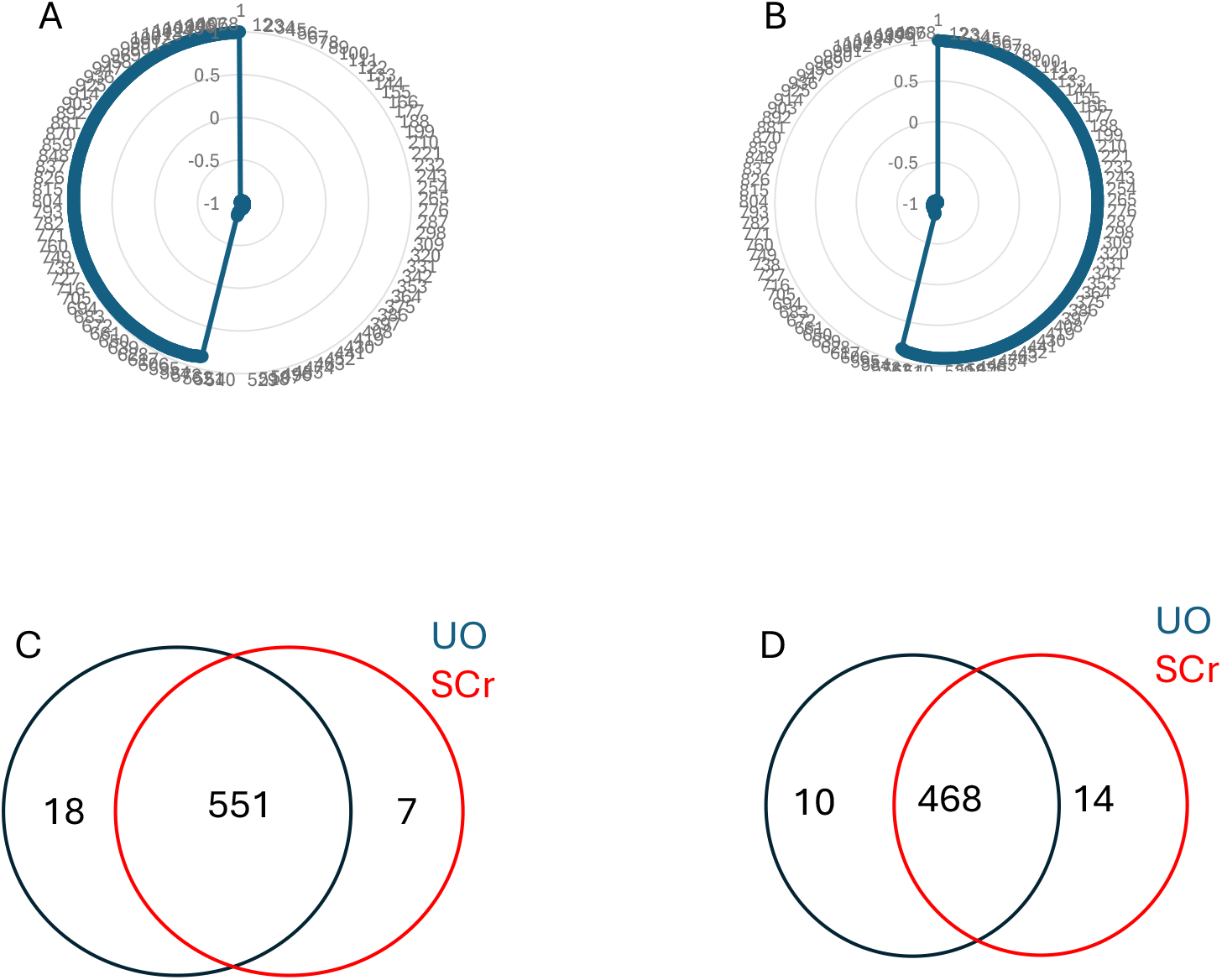
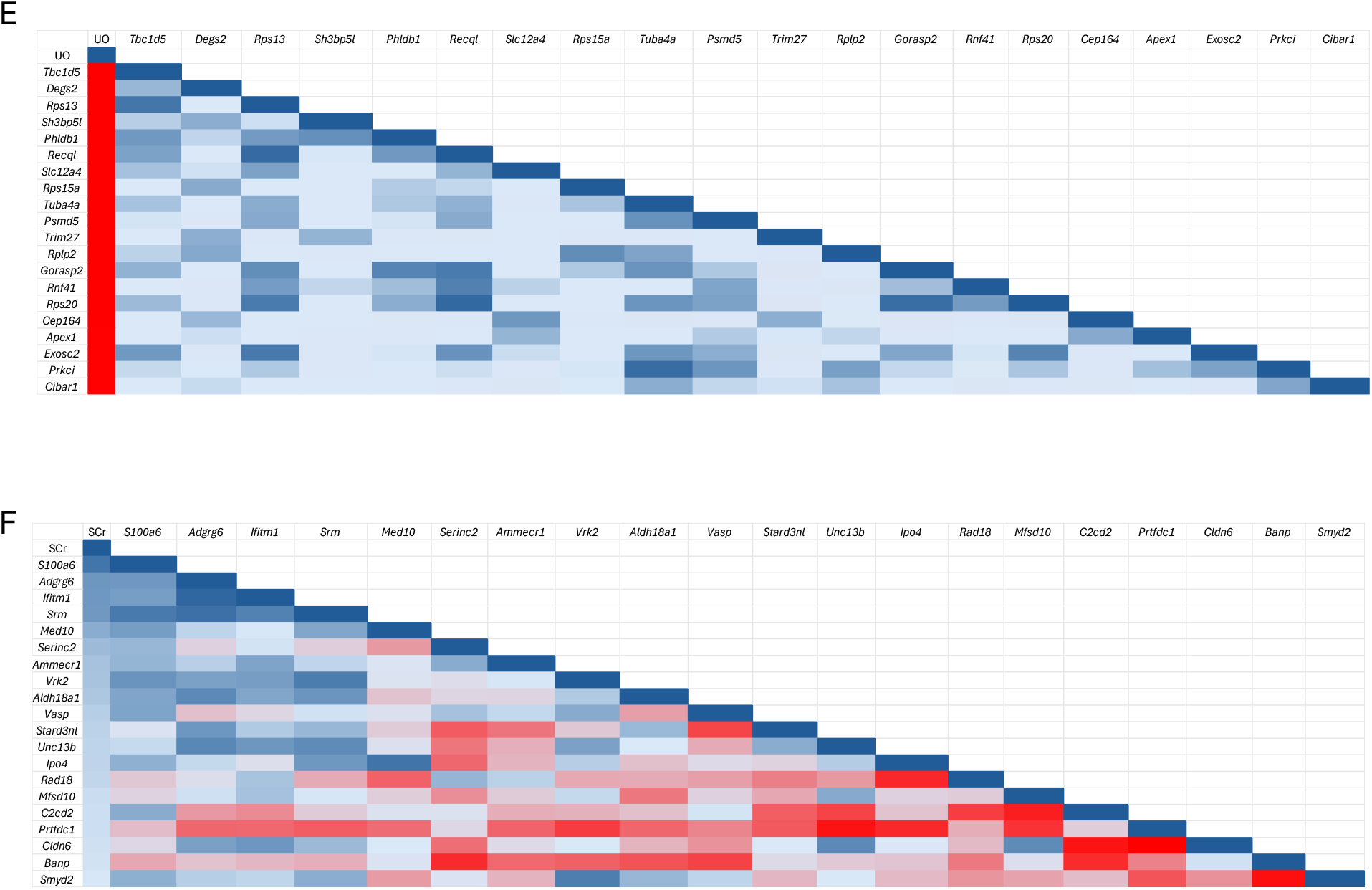
DEGs and Functional Correlations. Correlation (p<0.01) plots of DEGs vs. UO (A) and DEGs vs. SCr (B). The set of upregulated (C) and downregulated (D) genes correlating with UO *and* SCr shared a relatively high magnitude of similarity. The strength of the association (r) between DEGs correlating with UO (E, correlations for top 20 genes shown) was dissimilar to the strength of the association for DEGs correlating with SCr(F, correlations for top 20 genes shown).

To identify the pathway activation signature associated with oliguric AKI, up- and down regulated genes correlating with UO and SCr were submitted to GO biological process overrepresentation analysis. No biological pathway was associated with this set of genes. Next, the set of upregulated DEGs correlating with UO and SCr was submitted to overrepresentation analysis. The GO biological processes associated with this set are shown in Table 1 (> 5-fold enrichment only. Interestingly, this model of oliguric AKI was associated with a >44-fold enrichment in positive regulation of platelet derived-growth factor receptor (PDGFR)ß signaling pathway (Figures 4A and 4B). Three genes, src proto-oncogene tyrosine-protein kinase (*src*), Huntingtin interacting protein 1 related (*hip1r*) and Huntingtin interacting protein 1 (*hip1*) drove PDGFRß signaling. Activation of the PDGFRß signaling pathway is involved in transition of AKI to chronic kidney disease (CKD)^8,9^, a phenomenon involving macrophages^21^ and activated tubular epithelial cells^22^. Seeding of *pdgfrb, src, hip1r* and *hip1* into HumanBase revealed that each of these genes are relatively robustly expressed in the kidney (Figure 5A) and exhibit a relatively robust interactome amongst in macrophages (Figure 5A) and tubular epithelial cells (Figure 5B).

**Table 1.**
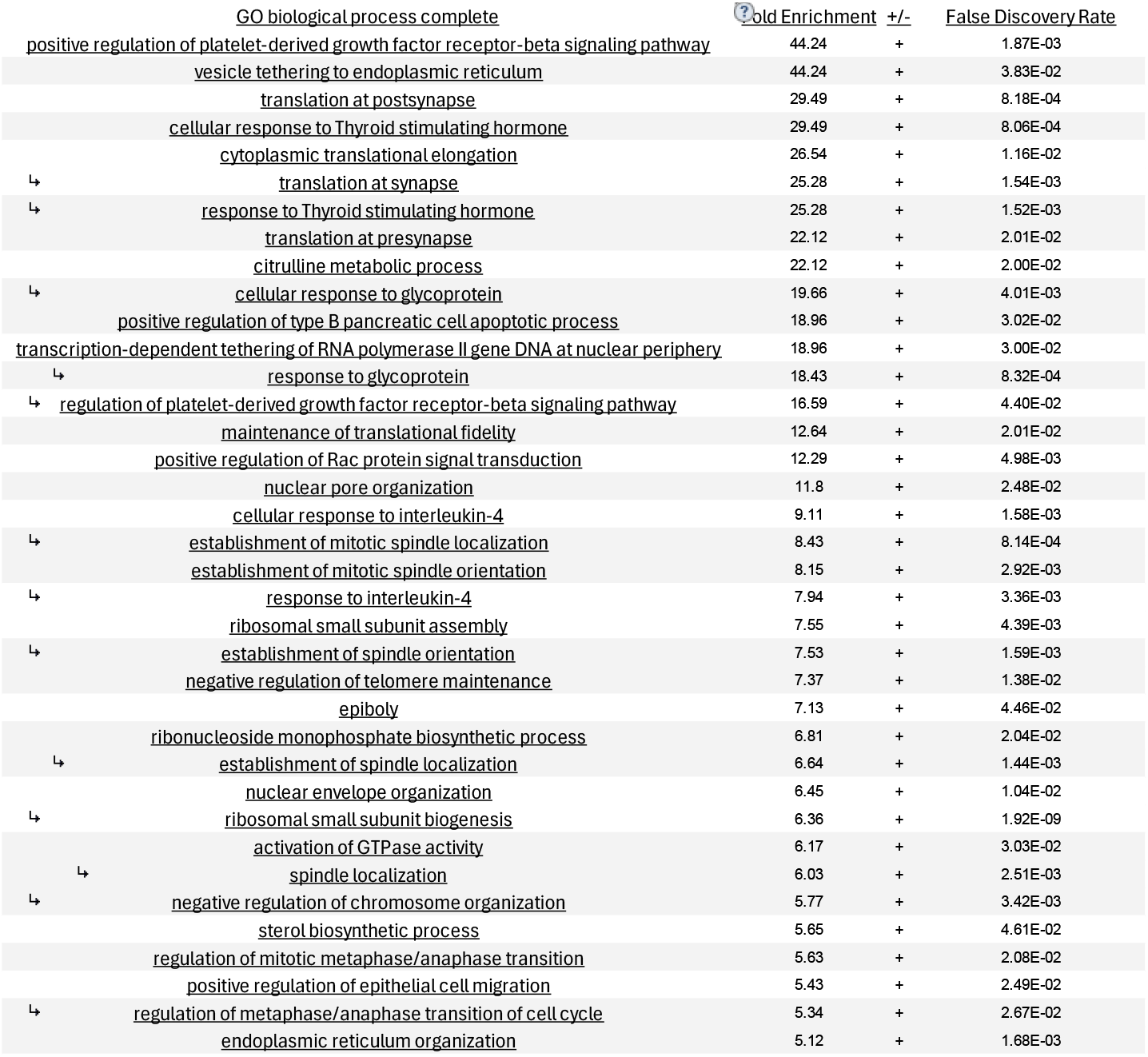
GO biological processes associated with our model of oliguric AKI.

|  | GO biological process complete | old Enrichment | +/- | False Discovery Rate |
| --- | --- | --- | --- | --- |
|  | <u>positive regulation of platelet-derived growth factor receptor-beta signaling pathway</u> | 44.24 | + | 1.87E-03 |
|  | <u>vesicle tethering to endoplasmic reticulum</u> | 44.24 | + | 3.83E-02 |
|  | <u>translation at postsynapse</u> | 29.49 | + | 8.18E-04 |
|  | <u>cellular response to Thyroid stimulating hormone</u> | 29.49 | + | 8.06E-04 |
|  | <u>cytoplasmic translational elongation</u> | 26.54 | + | 1.16E-02 |
| ↳ | <u>translation at synapse</u> | 25.28 | + | 1.54E-03 |
| ↳ | <u>response to Thyroid stimulating hormone</u> | 25.28 | + | 1.52E-03 |
|  | <u>translation at presynapse</u> | 22.12 | + | 2.01E-02 |
|  | <u>citrulline metabolic process</u> | 22.12 | + | 2.00E-02 |
| ↳ | <u>cellular response to glycoprotein</u> | 19.66 | + | 4.01E-03 |
|  | <u>positive regulation of type B pancreatic cell apoptotic process</u> | 18.96 | + | 3.02E-02 |
|  | <u>transcription-dependent tethering of RNA polymerase II gene DNA at nuclear periphery</u> | 18.96 | + | 3.00E-02 |
| ↳ | <u>response to glycoprotein</u> | 18.43 | + | 8.32E-04 |
| ↳ | <u>regulation of platelet-derived growth factor receptor-beta signaling pathway</u> | 16.59 | + | 4.40E-02 |
|  | <u>maintenance of translational fidelity</u> | 12.64 | + | 2.01E-02 |
|  | <u>positive regulation of Rac protein signal transduction</u> | 12.29 | + | 4.98E-03 |
|  | <u>nuclear pore organization</u> | 11.8 | + | 2.48E-02 |
|  | <u>cellular response to interleukin-4</u> | 9.11 | + | 1.58E-03 |
| ↳ | <u>establishment of mitotic spindle localization</u> | 8.43 | + | 8.14E-04 |
|  | <u>establishment of mitotic spindle orientation</u> | 8.15 | + | 2.92E-03 |
| ↳ | <u>response to interleukin-4</u> | 7.94 | + | 3.36E-03 |
|  | <u>ribosomal small subunit assembly</u> | 7.55 | + | 4.39E-03 |
| ↳ | <u>establishment of spindle orientation</u> | 7.53 | + | 1.59E-03 |
|  | <u>negative regulation of telomere maintenance</u> | 7.37 | + | 1.38E-02 |
|  | <u>epiboly</u> | 7.13 | + | 4.46E-02 |
|  | <u>ribonucleoside monophosphate biosynthetic process</u> | 6.81 | + | 2.04E-02 |
| ↳ | <u>establishment of spindle localization</u> | 6.64 | + | 1.44E-03 |
|  | <u>nuclear envelope organization</u> | 6.45 | + | 1.04E-02 |
| ↳ | <u>ribosomal small subunit biogenesis</u> | 6.36 | + | 1.92E-09 |
|  | <u>activation of GTPase activity</u> | 6.17 | + | 3.03E-02 |
| ↳ | <u>spindle localization</u> | 6.03 | + | 2.51E-03 |
| ↳ | <u>negative regulation of chromosome organization</u> | 5.77 | + | 3.42E-03 |
|  | <u>sterol biosynthetic process</u> | 5.65 | + | 4.61E-02 |
|  | <u>regulation of mitotic metaphase/anaphase transition</u> | 5.63 | + | 2.08E-02 |
|  | <u>positive regulation of epithelial cell migration</u> | 5.43 | + | 2.49E-02 |
| ↳ | <u>regulation of metaphase/anaphase transition of cell cycle</u> | 5.34 | + | 2.67E-02 |
|  | <u>endoplasmic reticulum organization</u> | 5.12 | + | 1.68E-03 |

**Figure 4.**
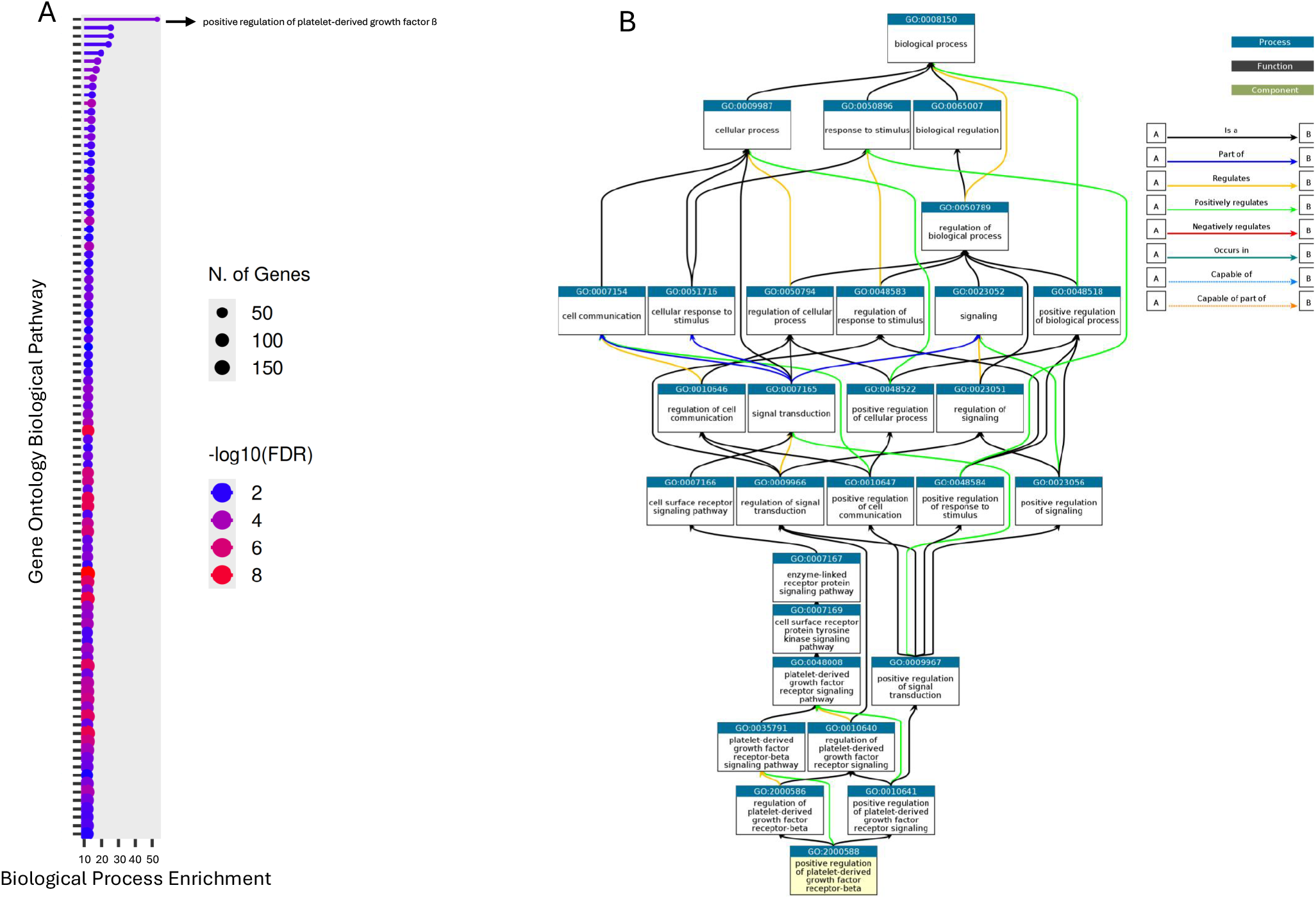
Activation of the PDGFRß Pathway in Oliguric AKI. (A) The PDGFRß pathway exhibited a fold-enrichment >44 in this model of oliguric AKI. (B) Ancestor chart for positive regulation of the PDGFRß pathway.

**Figure 5.**
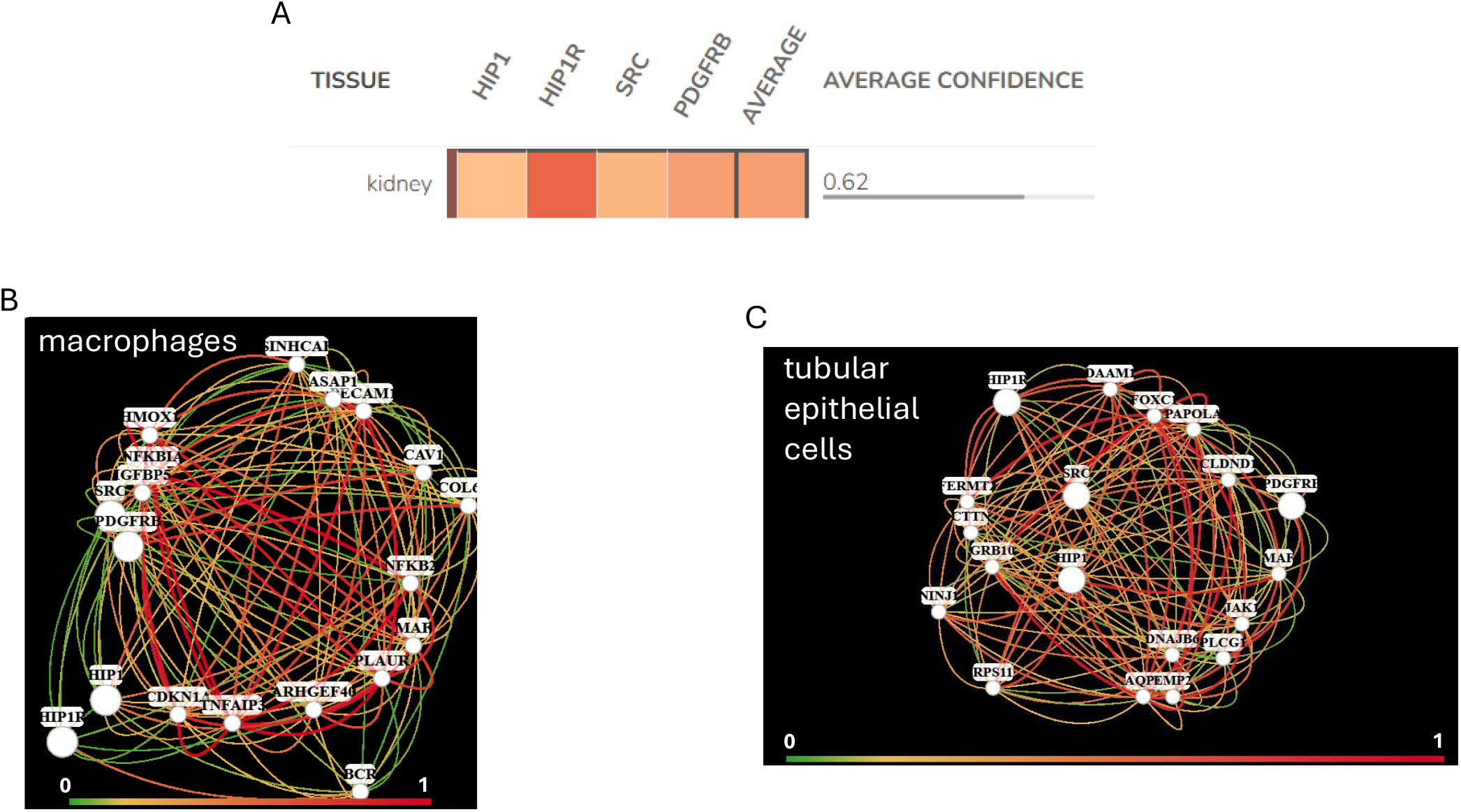
The *src, hip1* and hip1r Interactome. (A) Genes *src, hip1* and *hip1r* are expressed by the kidney and form relatively robust interactomes in macrophages (B) and tubukar epithelial cells (C).

## Discussion

In the present study we identified the pathway activation signature in a model of oliguric AKI. The activation signature includes positive regulation of the PDGFRß signaling pathway driven by upregulation of *src, hip1*, and *hip1r*. These data not only suggest that oliguric AKI may be associated with activation of a pathway leading to CKD but also provide a novel array of targets to mitigate sequalae of injury.

Annually affecting millions worldwide^2^, AKI can occur both outside and in hospital with its severity staged by the Kidney Disease: Improving Global Outcomes for AKI guidelines based on SCr increases or UO declines^23^. Outcomes in patients with AKI are polarizing and include an AKI to CKD transition triggered by incomplete or maladaptive kidney repair, leading to persistent inflammation, fibrosis, and a progressive reduction in filtration^7,24^. Oliguric AKI accompanied or AKI accompanied by a severe reduction in UO can be associated with outcomes distinct from AKI. Indeed, large cohort studies have demonstrated distinct and independent contributions to mortality and need for RRT with oliguric AKI^11-13^. In a model of transient oliguric AKI, Matsushita *et al*.^25^ reported a transition to CKD evidenced by a loss in glomerular filtration rate, proteinuria and increased kidney fibrosis.

In the present study we sought to derive the molecular pathway activation signature associated with oliguric AKI. The data queried were from a model of rat 45-minute unilateral kidney ischemia with 24 hours reperfusion1^4,15^. The contralateral kidney was removed at reperfusion. This model was accompanied by oliguria and an excursion in SCr. Bulk transcriptomic analysis revealed hallmark upregulation in *havcr1* which codes for KIM-1, and *lcn2* which codes for NGAL. Indeed, upregulation of these genes in the setting of AKI have been previously reported with their secreted gene products serving as biomarkers in AKI^19^. We used Boolean logic coupled with overrepresentation analysis to identify biological pathways associated with this model of oliguric AKI. The Boolean AND gate was used to first identify kidney DEGs associated with UO and SCr. The sets of downregulated and upregulated genes were submitted to overrepresentation analysis, a statistical method that determines if a specific set of genes is found more often in a particular pathway than would be expected by random chance^26^. A set of 551 upregulated genes and the associated GO biological processes formed the pathway activation signature in this model of oliguric AKI.

The most striking feature within this pathway activation signature was positive regulation of the PDGFRß signaling pathway. Indeed, this signaling pathway was dominant exhibiting >44-fold enrichment relative to background. The 3 upregulated genes, *src, hip1* and *hip1r*, driving this pathway are expressed in kidney tissue and form relatively robust interactomes with *pdgfrb* in both macrophages and activated tubular epithelium. These findings are of significance in the AKI-CKD transition, a potential sequel to oliguric AKI. The PDGRß signaling pathway plays a profibrotic role in multiple adult organs including the kidney mediating excessive extracellular matrix deposition^27,28^. Cytoplasmic tyrosine kinase SRC enhances PDGF signaling. The binding of PDGF to its PDGFRß leads to the recruitment and activation of SRC which in turn directly binds to activated PDGFRß, further amplifying the downstream signal^29^. HIP1, HIP1R, and PDGFRß are interconnected proteins with their interplay stabilizing PDGFRß on the cell surface and extending receptor half-life. Together, they regulate the strength and duration of the cellular response to PDGF^30^ allowing for a stronger and more sustained signaling cascade downstream of PDGFR and SRC. Expression of *src, hip1* and *hip1r* by macrophages and tubular epithelium is also significant. Pro-inflammatory macrophages and tubular epithelial cells engage in a dynamic crosstalk that drives maladaptive repair, inflammation, and fibrosis^21,22,24^. Indeed, antagonists to PDGFRß^31,32^ or SRC^33^ inhibited fibrotic processes in various organs, including the liver, lungs, and kidneys; the knockdown of HIP1R reduced fibroblast-like synoviocytes invasiveness and migration^34^, highlighting its role at least in inflammatory joint damage.

In summary, the pathway activation signature in oliguric AKI includes PDGFRß signaling driven by an *src-hip1-hip1r* axis, which may form, in part, the basis for AKI to CKD transition.

## Conclusion

Mechanistic insights should illuminate therapies. The pathway activation signature in oliguric AKI informs not only the sequel to injury but also an array of targets to mitigate the potential transition to kidney fibrosis and CKD.

## Supporting information

Fgure 3

